# Comparison of Non-linear and Linear Models of Single Channel EEG in patients and normal subjects

**DOI:** 10.1101/702605

**Authors:** Gu ZhuoJun, Huang ZhiQiang, Zhu Xiao, Shi ShenXun

## Abstract

This article examines the possibility of using non-linear models(Support Vector Regression) to model the single channel EEG signals from psychiatric patients and a group of normal participants, to predict psychology trait ratings, like attention, anxiety, alertness, fatigue, sleepiness and depression. It used linear models as benchmarks, and the results showed non-linear models outperformed the benchmarks, as well as more advanced linear methods, like principle component regression. It is thus concluded that using single channel in practical situations to monitor these traits would be possible.

## 1. Introduction

In recent years psychosomatic diseases from Kindergarten to Grade 12 has been requiring a lot of attention. Continuous monitoring of certain psychological traits are key issues in these fields. Among them, attention, anxiety, alert, fatigue, sleepiness and depression need to be examined carefully.

In psychology and medicine, researchers normally are using testing scales to measure these examined traits. However, in order to start this examination, a great number of items are needed for subjects to fill in. This process is time consuming and thus creates barriers for application. Some scales containing fewer items are developed in order to solve this issue, for example, Patient Health Questionnaire 9 (PHQ-9) for depression, and Generalized Anxiety Disorder scale 8 (GAD-8) for anxiety, which are common in clinical situations. But traits still cannot be assessed in real time.

Scales are also suffered from biases and cross-cultural validity due to the subjective nature of items. In practice, items are needed to be examined by domain experts to check their content validity. If it is used in a different culture, cross cultural validity becomes a notable issue. Thus, researchers need more objective tools to determine what can and cannot be used, a good example of a tool is electroencephalogram (EEG).

Both in research and practice, a multi-channel EEG device plays a major role. The range of numbers of typical EEG channels is from 32 to 256, according to the 10-20 EEG system standards^[1]^(Jaakko Malmivuo et al, 1995). The EEG needs lots of time to prepare before starting the recording^[2]^(Degabriele at al, 2008). The large number of electrodes also contributes to the high cost of the device. So for convenience reasons, reducing electrodes is the main concern.

Another major concern about EEG convenience is the nature of electrodes. Electrodes of most devices are wet, which require extra gel to increase the conductance. It hampers EEG going out from the lab^[3]^(Mathewson et al, 2017). An alternative choice would be the dry electrode, which is free from gel use.

Therefore, an ideal EEG device might contain 1 dry electrode attached to the location FP1 in the left side of the forehead of the person being tested, and using location FPZ as the ground electrode, which has already been widely used in EEG researches^[4][5]^(Johnstone et al, 2012; Rogers et al, 2016) with commercially available products, like Neurosky mobile EEG set.

However, researchers can measure traits using scales directly. On the contrary, one cannot use EEG in the same way. And rare researches have been done to investigate the relationship between psychological traits and EEG measures, especially using single channel device. In psychology literatures, despite of the availability of 1 channel EEG device, multi-channel devices were still used in most researches, like attention^[6]^ (Gevins et al, 1997), anxiety and depression^[7]^(Smit et al, 2007), alert and fatigue^[8]^(Tran et al, 2008), sleepiness^[9]^(Herron et al, 2014). Thus, there are needs to explore relationships between traits and single channel EEG. But what we face here is the problem of non-linearity.

Statistics has been widely accepted in EEG research. Most articles use linear models, like generalized linear model^[10]^ (Redelico et al, 2017) or general linear model^[11]^ (Moeller et al, 2011), to analyze the relationship between traits and EEG measures. The shortcoming of these models is excluding the non-linear relationships, but brain is a non-linear system^[12]^ (Andrzejak et al, 2001), so there might be possibilities that non-linear models may outperform linear models. For convenience, the difference between linearity and non-linearity is illustrated in the following equation 1.1:

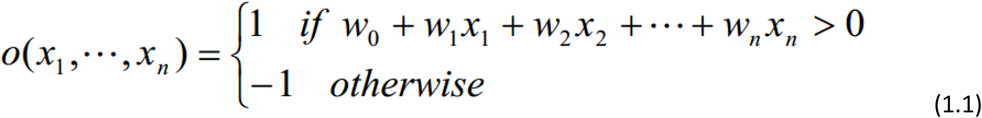

This is an artificial neural network (ANN) model called “perceptron” ^[13]^ (Mitchell, 2003). It calculates the result of a linear equation of vector (x_1_, …, x_n_), where w_n_ refers to weights of the variable x_n_ similar to beta in linear regressions. If the result is larger than 0, it outputs 1, otherwise it gives −1. ANN is in nature a non-linear model, while it can still contain a linear part.

Support Vector Machine (SVM)^[14]^(Alex et al, 2004) as a typical non-linear model in machine learning, has also been widely used in EEG, like in sleep^[15]^(Lajnef et al, 2015), motor tasks^[16]^(Zhou et al, 2007), epilepsy^[17]^(Temko et al, 2011), source localization^[18]^(Besserve et al, 2011) and mental imagery^[19]^(Xu et al, 2009). Current researches mainly use the classification method, rather than the regression method. While both generalized and general linear models are regression based methods, SVM does have a regression model^[20]^ (Pedregosa et al, 2011), which is called “Support Vector Regression”(SVG). Since the brain is non-linear, regression-based non-linear SVM may beat the generalized linear regression method, i.e., principle component Ordinal Logistic Regression. And we expect it would be also valid for linear models dealing with co-linearity, which would be also used in our analysis, including Ridge^[21]^(Tutz et al, 2005) and Partial Least Squares(PLS) Regression^[22]^(Geladi et al, 1986) (or cross decomposition regression in Python).

Unlike statistical regression methods, SVG is in nature a process of machine learning. The process needs optimization methods to find a better solution^[23]^(Mitchell, 2003). Among all optimization methods, Dual Annealing^[24]^(Xiang et al, 2013) and modified Powell method^[25]^(Zhang et al, 2013) would be used. Dual Annealing firstly would find a near-optimal solution using simulated annealing algorithm, then it incorporates modified Powell method to look for a local optimal solution. Compared to Maximum Likelihood Estimation (MLE) method used in the statistical regression, these methods are superior since they do not require the estimation function to be differentiable. We implement the optimization method to search for a better model with the larger R^2^.

The main idea of this article is to use SVG model to analyze the relationship between multiple traits’ ratings and single channel EEG data. It is expected that single channel EEG can be used to predict these traits, and non-linear models (SVG) would be more advantageous than linear ones.

## 2. Methods

### 2.1 Study design and participant

134 Chinese people were invited to participate in two groups which are: the patient and the normal group. 80 participants in the patient group were diagnosed with psychiatric disorders by psychiatrists. 54 participants in the normal group are employees from the company.

The age of patients ranges from 15 to 67 (34.49 in average and its stand deviation: 13). Among them, 32 were male and 48 were female. They were tested in the Department of Psychiatry, HuaShan Hospital, East Branch in Shanghai.

The age of the participants in the normal group is within the range of 21 to 53 (30.12 in average and its stand deviation: 9.08). Among them, 26 were male and 28 were female. They were tested in a separated conference rooms in the company’s building.

All patients were informed about their rights based on certain research ethics of the hospital. All normal people were asked to help to participate in the research, as an aid in the development of the EEG device.

All participants signed the informed consent before the study began. All participants claimed they had not taken psychiatric drugs, drunken coffee or alcohol in the last 24 hours before the study.

### 2.2 Protocol and measurements

Before the study patients completed two scales administered by psychiatrists in the hospital: PHQ-9^[25]^(Kroenke et al, 2001) for depression and GAD-7^[27]^(Lowe et al, 2008) for anxiety. Then they signed the informed consent. Normal group started to fill in the informed consent directly.

A single-channel EEG device (developed by Shanghai HuiCheng Science & Education Equipment Company) was worn by participants to collect EEG data in FP1 channel of 10-20 system. And the device used FPz channel as the ground electrode. EEG was recorded at a sampling rate of 500Hz.

Before they wore it, the electrodes were sterilized by alcohol. After it was cleansed they were asked to put the device on, assuring the electrodes were correctly positioned. The person being tested is seated in front of a desk, where a computer screen, a keyboard and a mouse were, along with an A4 paper with a black cross symbol in the center was beside the screen.

Participants performed tasks that each lasted 4-minutes. In the first 2 minutes, they kept their eyes focused on the cross on the paper. And then the screen showed a Schulte Table^[28]^(Sosin et al, 2016) with 8 × 8 cells showing numbers from 1 to 64 in a random order. For the next 2 minutes, the testie used the mouse to click the numbers starting from 1.

After the task, they were asked to rate their current levels of attention, anxiety, alertness, fatigue, sleepiness and on a 9-points (1 to 9) semantic differential scale. In the scale, if the grade is higher for attention and alertness, it means participants have higher levels of attention and alertness. Other traits have the opposite relationships between grades and levels. An experimenter guided each participant undergoing the whole test, and wrote down any behaviors and their time disrupting the EEG recording, like body moving or question asking.

### 2.3 EEG preparation and artifact removal

The EEG device had a built-in denoising and artifact processing pipeline, including low and high band-pass filters, 50Hz band-pass filter, filters used to process artifacts like head movements and eye movements. The time-frequency decomposition method was wavelet transform[29]^[29]^(Li et al, 2009). And the mother wavelet was Daubechies order 4^[30]^(Adeli et al, 2003). It output 4 minutes raw EEG data of each participant of FP1 channel, a total of 240 points of EEG raw data, containing powers of Delta, Theta, Low Alpha, High Alpha, Low Beta, High Beta, and Gamma.

The first 3 minutes were discarded. In the last 1 minute data, time points of pre-recorded disrupting behaviors were also excluded. Grubbs’ test was taken to detect outliers, in which the significance criterion of the test was 0.05. After outlier removal, natural logarithmic values of the 1m average were obtained. Then we used Z value of the data for further analysis, ensuring the data were in the same range.

### 2.4 Data analysis

SPSS (version 23) and Python machine learning package scikit-learn^[20]^ (or Sklearn, version 0.20.2) were used to conduct the data analysis, while Python scientific computing package SciPy (version 1.2.1) was used to prioritize the linear and non-linear models’ hyper-parameters.

The consistencies between psychiatric scales and ratings were checked in SPSS. We calculated Spearman non-parametric correlation rho between depression rating and PHQ-9, and between anxiety rating and GAD-7.

For traits: attention, anxiety, alertness, fatigue, sleepiness, the following steps were conducted.

Firstly, an exploratory principle component analysis was done in SPSS, in which the extraction method was the principal component, to help determine the number of latent factors and deal with co-linearity among EEG frequency bands for linear modeling. Then we calculate the factor score based on the results.

Then generalized linear model of Ordinal Logistic was done in SPSS between the factor and the trait score. This analysis is used as a benchmark. We record Nagelkerke’s R^2^ in SPSS as the benchmark index. This is because it has a range of 0 to 1^[31]^(Nagelkerke, 1991).

Next we use both linear and non-linear models in Sklearn to explore the relationships between EEG frequency bands and subjective ratings. PLS regression and Ridge were used in linear modeling. As for non-linear modeling, NuSVM with the linear kernel and radial basis function (RBF) kernel^[32]^(Chang et al, 2011) were used.

Each model had its hyper-parameters to be optimized according to certain validation measures. We chose to optimize the most relevant parameters. PLS regression had 1, “n_components”, derived from the factor analysis in SPSS. So it was calculated manually and was excluded from the prioritization process.

During the process, hyper-parameters^1^ were tuned. Ridge had 1, alpha(range from 0.1 to 50). NuSVM with the linear kernel has 1, nu(range: from 0.01 to 0.99) and C(range from 0.5 to 5). NuSVM with the RBF kernel had 2, nu(range: from 0.01 to 0.99), C(range from 0.5 to 5) and gamma(range: from 0.1 to 2). Detailed meanings of each hyper-parameter are accessible via the Sklearn online documentation. All other hyper-parameters were set default.

The measure to be optimized was a composite of the model bias and the model variance.

For evaluating the bias, we incorporated a method called “Leave-One-Out” stragegy^[33]^(Peter et al., 2014), meaning leaving one sample in the testing set and using all other samples to train the model until every sample had been left once in the testing set, and finally calculating the difference measure like R^2^ between the predicted values and the actual values, i.e., 1 minus the quotient which squared sum of error divides total sum of error (STD), which the range was from negative infinity to 1. The reason using R^2^ rather than the traditional mean standard deviation here is due to the ease of explanation.

To evaluate the variance, the standard deviation (STD) of predicted values of certain hyper-parameters were calculated.

We constructed the validation measure for non-linear models by using −0.8 multiplied leave one out R squared plus −0.1 multiplied model R^2^, adding 0.05 multiplied STD, and adding 0.05 multiplied Nu, meaning the quantity of support vectors in non-linear models defined the model complexity. For linear models, the STD weight was changed to 0.1 and Nu parameter was excluded. Then we used an optimization algorithm called “dual annealing” in Scipy optimize module, which was to find the minimum of this measure. Dual annealing was a two-step algorithm. First it ran the simulated annealing, which simulates the molecule cooling process in nature, to find a good solution near the global minimum, then it used the modified Powell method to find the ideal minimum as the near-perfect solution. Due to the stochastic aspect of the algorithm^[34]^(Mullen, 2014), totally we ran 10 epochs. In each epoch, we started with random initial values of hyper-parameters. For the first 5 epochs, we chose the hyper-parameter values based on the minimum of the validation measure. During the last 5 epochs, we started each epoch with the best value among the first 5 epochs to check whether any better values could be found.

The cross validation was run in Python 3.7 environment, using Pycharm Integrated Development Environment (IDE) version 2018.3.5 community edition.

Due to application reasons and stability of depression, another procedure was performed on it:

We filtered the patient group based on their digital medical records, locating patients who were diagnosed with depression disorder, depressed state or mania (depressed currently). In total there were 49 patients. Then we combined the normal group with the depression patient group, and calculated the EEG factor score of every participant based on a factor analysis using principle component extraction, where only 1 factor was extracted. Next we conducted Linear Discriminant Analysis and Categorical and Regression Tree (CRT) ^[35]^(Li, 2012)using EEG factor score as the predict variable to predict the patient-or-not dummy variable.

## 3. Results

### 3.1 Analysis of attention, anxiety, alertness, fatigue, sleepiness

#### 3.1.1 Factor analysis of EEG bands

**Table 3.1.1 - a.** KMO and Bartlett’s test of Spherity.

|  |  |  |
| --- | --- | --- |
| KMO |  | .866 |
| Bartlett's test of Spherity | Chi-square (Appro.) | 2473.675 |
|  | Degree of Freedom | 21 |
|  | Sig. | <0.001 |

**Table 3.1.1 - b.** Total Variance Explained.

| Factor | Initial eigen value |  |  | Squared sum of factor loadings |  |  |
| --- | --- | --- | --- | --- | --- | --- |
|  | Total | Variance percentage | Cumulative % | Total | Variance percentage | Cumulative % |
| 1 | 6.498 | 92.831 | 92.831 | 6.498 | 92.831 | 92.831 |
| 2 | .348 | 4.966 | 97.797 |  |  |  |
| 3 | .103 | 1.469 | 99.266 |  |  |  |
| 4 | .017 | .243 | 99.509 |  |  |  |
| 5 | .016 | .226 | 99.735 |  |  |  |
| 6 | .010 | .147 | 99.882 |  |  |  |
| 7 | .008 | .118 | 100.000 |  |  |  |

Extraction: Principle Component

The KMO and Bartlett’s test of Spherity in Table 3.2.1 - a were acceptable. And 1 factor was extracted, shown in Table 3.1.1 - b, deriving the common component of EEG using component scores. We used it as the predictor of ratings as the benchmark, using Ordinal Logistic Regression method, shown in section 3.1.2

#### 3.1.2 Result of models

##### a) Attention, Anxiety, Alertness, Fatigue

According to Table 3.1.2 – a1, RBF SVM model beat all other models in internal R^2^ metric, including non-linear model linear SVM. And it is the only model whose leave_one_out R^2^ is larger than zero, meaning it was better than using the mean rating to predict. The internal R^2^ metric of RBF SVM is also larger than that of Ordinal logistic benchmark.

**Table 3.1.2 - a1^2^.** attention results.

| regression method | linear/non-linear | internal R<sup>2</sup> | leave_one_out R<sup>2</sup> | parameter 1 | parameter 2 | parameter 3 |
| --- | --- | --- | --- | --- | --- | --- |
| Ordinal logistic | benchmark | 0.003 | / | / | / | / |
| PLS | linear | 0.02 | -0.02 | number of components: 1 | / | / |
| Ridge | linear | 0.02 | -0.03 | alpha = 50 | / | / |
| linear SVM | non-linear | -0.02 | -0.02 | Nu = 0.01 | C = 0.5 | / |
| RBF SVM | non-linear | 0.18 | 0.02 | Nu = 0.06 | C = 2.22 | gamma = 1.12 |

In Table 3.1.2 - a2, the result is the same as in attention section.

**Table 3.1.2 - a2.**
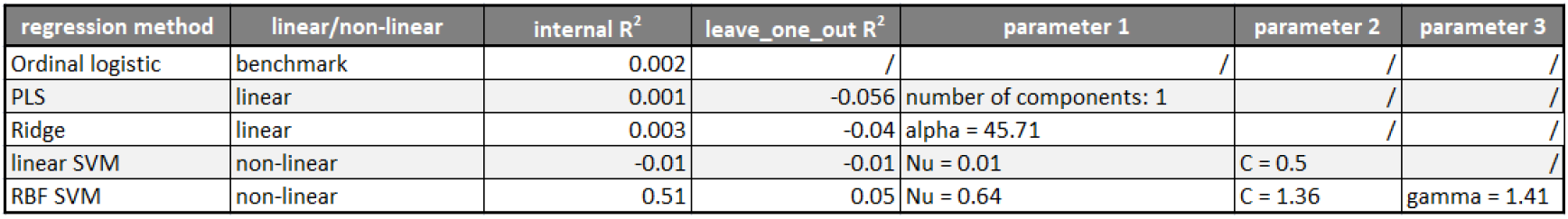
anxiety results.

| regression method | linear/non-linear | internal R<sup>2</sup> | leave_one_out R<sup>2</sup> | parameter 1 | parameter 2 | parameter 3 |
| --- | --- | --- | --- | --- | --- | --- |
| Ordinal logistic | benchmark | 0.002 | / | / | / | / |
| PLS | linear | 0.001 | -0.056 | number of components: 1 | / | / |
| Ridge | linear | 0.003 | -0.04 | alpha = 45.71 | / | / |
| linear SVM | non-linear | -0.01 | -0.01 | Nu = 0.01 | C = 0.5 | / |
| RBF SVM | non-linear | 0.51 | 0.05 | Nu = 0.64 | C = 1.36 | gamma = 1.41 |

In Table 3.1.2 - a3, Alertness results have verified the results in attention section again.

**Table 3.1.2 - a3.** alertness results.

| regression method | linear/non-linear | internal R<sup>2</sup> | leave_one_out R<sup>2</sup> | parameter 1 | parameter 2 | parameter 3 |
| --- | --- | --- | --- | --- | --- | --- |
| Ordinal logistic | benchmark | 0.003 | / | / | / | / |
| PLS | linear | 0.01 | -0.03 | number of components: 1 | / | / |
| Ridge | linear | 0.008 | -0.039 | alpha = 43.9 | / | / |
| linear SVM | non-linear | 0.001 | -0.011 | Nu = 0.18 | C = 0.5 | / |
| RBF SVM | non-linear | 0.21 | 0.061 | Nu = 0.79 | C = 5 | gamma = 0.1 |

In Table 3.1.2 - a4, the same results came up again.

**Table 3.1.2 - a4.**
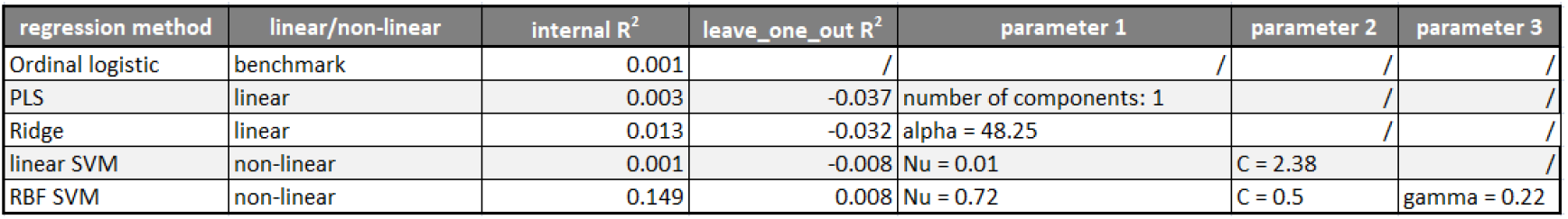
fatigue results.

##### b) Sleepiness

In Table 3.1.2 - b, the results of sleepiness differed. Linear SVM appeared to be better than RBF SVM, and both of their leave_one_out R^2^ was larger than 0. But all non-linear models still performed better than linear models, including the benchmark.

**Table 3.1.2 - b.**
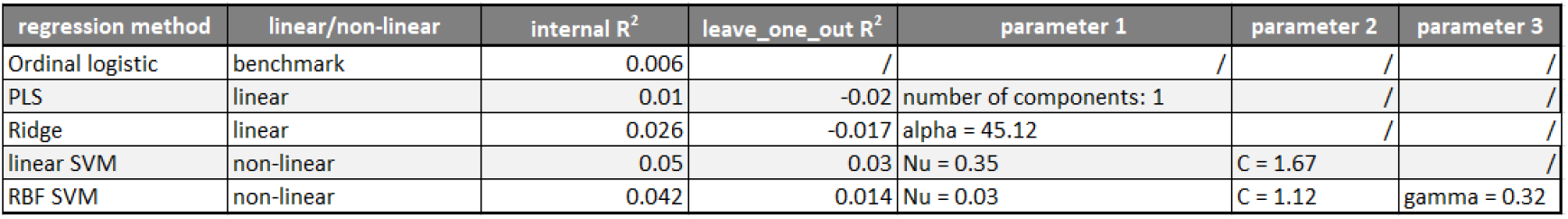
sleepiness results.

### 3.2 Analysis of depression

Unlike other trait ratings, SVM models of depression does not performed well where leave_one_out R2 of the optimal model was smaller than 0. So here we adopted decision tree model which was also used to model nonlinearity^[36]^(Mitchell, 2003).

An initial factor analysis of 103 participants extracted only 1 component which explained 92.605% of total variance, where KMO was 0.866 and P value of Bartlett’s test of Spherity was smaller than 0.001. Further analysis of the EEG factor score was taken.

To better understand the difference between linear and non-linear models, a method called confusion matrix was used. The confusion matrix of linear and non-linear models between the patient-or-not dummy variable and the EEG factor score were as follows.

**Table 3.2 - a.** Confusion Matrix a. linear discriminant analysis

|  | predict |  |  |
| --- | --- | --- | --- |
|  | 0 | 1 | Percent of correct prediction |
| raw,1 = patient |  |  |  |
| 0 | 37 | 17 | 68.5% |
| 1 | 32 | 17 | 34.7% |
| Average percent of correct prediction |  |  | 52.4% |

**Table 3.2 - b.** Confusion Matrix b. CRT

|  | predict |  |  |
| --- | --- | --- | --- |
|  | 0 | 1 | Percent of correct prediction |
| raw,1 = patient |  |  |  |
| 0 | 51 | 3 | 94.4% |
| 1 | 39 | 10 | 20.4% |
| Average percent of correct prediction |  |  | 59.2% |

The non-linear CRT model performs (59.2%) better than that of linear dicriminant analysis(52.4%).

### 3.3 Consistencies between Psychiatric Scales and ratings

**Table 3.3.** As shown in Table 3.3, anxiety rating is significantly correlated with GAD-7. It is similar between depression rating and PHQ-9.

|  |  | anxiety | depression | PHQ-9 | GAD-7 |
| --- | --- | --- | --- | --- | --- |
| anxiety | Pearson R | 1 | .268* | .099 | .236* |
|  | Sig.(two tail) | / | .016 | .384 | .035 |
|  | Sample | 80 | 80 | 80 | 80 |
| depression | Pearson R | .268* | 1 | .484** | .357** |
|  | Sig.(two tail) | .016 | / | < .001 | .001 |
|  | Sample | 80 | 80 | 80 | 80 |
| PHQ-9 | Pearson R | .099 | .484** | 1 | .740** |
|  | Sig.(two tail) | .384 | < .001 | / | < 0.001 |
|  | Sample | 80 | 80 | 80 | 80 |
| GAD-7 | Pearson R | .236* | .357** | .740** | 1 |
|  | Sig.(two tail) | .035 | .001 | < .001 | / |
|  | Sample | 80 | 80 | 80 | 80 |

\*. 0.05
\*\*. 0.01

## 4. Discussion

### 4.1 Validity of single-channel EEG prediction

Results using SVM between the single-channel EEG band and the trait scores showed ratings could be predicted by these band powers in the forehead. This is probable at least for attention, anxiety, alertness, fatigue and sleepiness. For categorizing the levels of depression, it should also help, but may not be the most accurate.

Ratings have been proven to measure how the mental trait varies since the early days of psychology, but it is still subject to drawbacks of wording issues, like content validity and inter-cultural validity, which have not been improved yet. When researchers want to implement ratings or scales in different groups or environments, items are always needing to be modified. Due to the objectivity of EEG signal, it sheds light on solving the above issue.

Another problem with EEG is the high cost of multi-channels and gel using. The feasibility of a single-channel EEG with the dry electrode minimizes it. In the data collection phase of our research, the preparation time was always less than multiple seconds. This allows wider usage of EEG devices, especially in the out-of-lab, commercial context.

It is expected that single-channel EEG has the potential to predict more trait ratings. By using single-channel EEG and more powerful modeling methods, more and more objective measures of traits could be obtained.

### 4.2 About trait ratings’ usage

EEG successfully predicts all the trait ratings. In current research, depression, anxiety, attention have validity criterion. They are PHQ-9 for depression, GAD-7 for anxiety, and Schulte-Grid task for attention, where the former two have significant correlation. Due to the successful prediction, the current research provides evidence that single-channel EEG could be used in contexts where it needs measures of the 3 mental traits.

Due to time and budget, traits like fatigue, alertness, and sleepiness do not have validity criterion in this research. The success of EEG prediction here suggests that these three might be used in out-of-lab contexts, which may require further research with their validity criterion.

### 4.3 Advantageous non-linearity modeling between EEG and trait scores

As expected, non-linear models of all traits performed better than linear models. The internal R^2^ had far better increases in RBF SVM, compared to the benchmark generalized linear regression.

Z-transformed rating is widely used, which is presumed as a continuum of mental traits in psychology and marketing, like the attitude rating scale, by increasing the number of samples. This research showed that SVM regression could be helpful in modeling their relationships, and their relationships are probably non-linear. This research may also remind us of the current limitations of scales used in psychology, indicating possible future directions of improvements.

### 4.4 Non-linearity in EEG

Past researches successfully used SVM to categorize mental status, like alert and fatigue^[8]^(Tran et al, 2008), sleepiness^[9]^(Herron et al, 2014). Though current research used SVM regression method, it is still non-linear in nature. So the result is consistent to past researches. This is another evidence that brain is a non-linear system^[12]^ (Andrzejak et al, 2001), which requires non-linear models to analyze.

Also, researchers have created non-linear metrics, like entropy measures, to reflect non-linearity in EEG. Although current research did not use these metrics, its use of common frequency bands is more commonly used in psychology and a wider community. It should be a sounder piece of evidence of EEG non-linearity than entropy measures.

## 5. Limitations and future work

The present article only used single channel EEG device, and the results need to be validated further in multi-channel devices. Also, the results were based on mathematical models which lack interpretability. More functional neuroimaging(fMRI) studies should be taken to investigate its physiology. This is especially true to explain to the difference of models between depression and other ratings. These results are only covering opening eyes, and whether they are similar in participants closing their eyes is not sure, because the alpha band would become smaller in these situations.

In the future, as stated above, due to the nonlinearity of the brain, it should help by aiding entropy measures into the modeling process. Further analysis should also investigate situations of closing eyes.

## 6. Conclusion

The current research showed a single-channel EEG with 1 dry electrode could be used to predict trait scores. And their relationships are non-linear. Due to the ease of single channel, in practical situations, multiple psychology traits could be monitored using this device.

## Footnotes

1 The ranges of hyper-parameters here were based on previous exploratory numerical experiments, where leave-one-out R^2^ fell into a reasonable range, like from −1 to 0.2.

2 Ordinal logistic used the factor score of all EEG bands as the predictor, while other models put all bands into the regression. (It was the same for other ratings.)

## References

[1]. Jaakko Malmivuo, Robert Plonsey. “Bioelectromagnetism – Principles and Applications of Bioelectric and Biomagnetic Fields”. Oxford University Press, 1995, pp. 365–374.

[2]. Racheal Degabriele, Jim Lagopoulos. “Techniques for effective EEG subject preparation”. Acta Neuropsychiatrica. 2008, 20: 218–219.

[3]. Kyle E. Mathewson, Tyler J. L. Harrisson and Sayeed A. D. Kizuk. “High and dry? Comparing active dry EEG electrodes to active and passive wet”. Psychophysiology. 2017, 54: 74–82.

[4]. Stuart J. Johnstone, R Blackman and Jason Bruggemann. “EEG from a single-channel dry-sensor recording device”. Clinical EEG and Neuroscience. 2012, 43: 112–120.

[5]. Jeffrey M. Rogers, Stuart J. Johnstone, Anna Aminov, James Donnelly and Peter H. Wilson. “Test-retest reliability of a single-channel, wireless EEG system”. International Journal of Psychophysiology. 2016, 106: 87–96.

[6]. Alan Gevins, Michael E. Smith, Linda McEvoy and Daphne Yu. “High-resolution EEG Mapping of Cortical Activation Related to Working Memory: Effects of Task Difficulty, Type of Processing, and Practice”. Cerebral Cortex. 1997, 7: 374–385.

[7]. D.J.A. Smit, D. Posthuma, D.I. Boomsma and E.J.C. De Geus. “The relation between frontal EEG asymmetry and the risk of anxiety and depression”. Biological Psychology. 2007, 74: 26–33.

[8]. Y. Tran, N. Wijesuryia, R.A. Thuraisingham, A. Craig, and H.T. Nguyen. “Increase in regularity and decrease in variability seen in electroencephalography (EEG) signals from alert to fatigue during a driving simulated task”. Engineering in Medicine and Biology Society, 2008. EMBS 2008. 30th Annual International Conference of the IEEE, 20–25 Aug. 2008.

[9]. Katherine Herron, Derk-Jan Dijk, Philip Dean, Ellen Seiss and Annette Sterr. “Quantitative Electroencephalography and Behavioural Correlates of Daytime Sleepiness in Chronic Stroke”. BioMed Research International. 2014.

[10]. Francisco O. Redelico, Francisco Traversaro, María del Carmen García, Walter Silva, Osvaldo A. Rosso and Marcelo Risk. “Classification of Normal and Pre-Ictal EEG Signals Using Permutation Entropies and a Generalized Linear Model as a Classifier”. Entropy. 2017, 19(2), 72.

[11]. Friederike Moeller, Pierre LeVan and Jean Gotman. “Independent Component Analysis (ICA) of Generalized Spike Wave Discharges in fMRI: Comparison with General Linear Model-Based EEG-fMRI”. Hum Brain Mapping. 2011, 32(2): 209–217.

[12]. Ralph G. Andrzejak, Klaus Lehnertz, Florian Mormann, Christoph Rieke, Peter David and Christian E. Elger. “Indications of non-linear deterministic and finite-dimensional structures in time series of brain electrical activity: Dependence on recording region and brain state”. PHYSICAL REVIEW E. 2001, 64.

[13]. Mitchell, T. M. “Machine Learning”. China Machine Press. 2003, p.87.

[14]. Alex J. Smola, Bernhard Schölkopf. “A Tutorial on Support Vector Regression”. Statistics and Computing. 2004, 14: 199–222.

[15]. Tarek Lajnef, Sahbi Chaibi, Perrine Ruby, Pierre-Emmanuel Aguera, Jean-Baptiste Eichenlaub, Mounir Samet, Abdennaceur Kachouri and Karim Jerbi. “Learning machines and sleeping brains: Automatic sleep stage classification using decision-tree multi-class support vector machines”. Journal of Neuroscience Methods. 2015, 250: 94–105.

[16]. S-M. Zhou, J.Q. Gan and F. Sepulveda. “Classifying Mental Tasks Based on Features of HigherOrder Statistics from EEG Signals in Brain-Computer Interface”. Information Sciences. 2007.

[17]. A. Temko, E. Thomas, W. Marnane, G. Lightbody and G.B. Boylan. “Performance assessment for EEG-based neonatal seizure detectors”. Clinical Neurophysiology. 2011, 122: 474–482.

[18]. Michel Besserve, Jacques Martinerie and Line Garnero. “Improving quantification of functional networks with EEG inverse problem: Evidence from a decoding point of view”. NeuroImage. 2011, 55: 1536–1547.

[19]. Qi Xu, Hui Zhou, Yongji Wang, Jian Huang. “Fuzzy support vector machine for classification of EEG signals using wavelet-based features”. Medical Engineering & Physics. 2009, 31: 858–865.

[20]. Fabian Pedregosa, Gaël Varoquaux, Alexandre Gramfort, Vincent Michel, Bertrand Thirion, Olivier Grisel, Mathieu Blondel, Peter Prettenhofer, Ron Weiss, Vincent Dubourg, Jake Vanderplas, Alexandre Passos, David Cournapeau, Matthieu Brucher, Matthieu Perrot and Édouard Duchesnay. “Scikit-learn: Machine Learning in Python”. Journal of Machine Learning Research. 2011, 12: 2825–2830.

[21]. Tutz G, Binder H. “Boosting ridge regression”. Computational Statistics and Data Analysis. 2007, 51(12):6044–6059.

[22]. Geladi P, Kowalski B R. “Partial Least-Squares Regression: A Tutorial”. Analytica Chimica Acta. 1986, 185(1):1–17.

[23]. Mitchell, T. M. “Machine Learning”. China Machine Press. 2003, p.267.

[24]. Xiang Y., Gubian S., Suomela B. and Hoeng Julia. “Generalized Simulated Annealing for Global Optimization: The GenSA Package.” The R Journal. 2013, 5(1): 13–28.

[25]. Zhang K. and Luo J. “Research on flatness errors evaluation based on artificial fish swarm algorithm and Powell method”. Int. J. Computing Science and Mathematics. 2013, 4(4):402–411.

[26]. Kroenke K., Spitzer R. L. and Williams J. B. W. “The PHQ-9 : Validity of a Brief Depression Severity Measure”. Journal of General Internal Medicine. 2001, 16(9):606–613.

[27]. Lowe B. and Schellberg D. “Validation and Standardization of the Generalized Anxiety Disorder Screener (GAD-7) in the General Population”. Medical Care. 2008, 46(3):266–274.

[28]. Sosin I., Chuev Y. and Goncharova O. “Problems of activation of attention psychophysiological functions and peripheral visual perception of traffic lights color analogues”. European Psychiartry. 2016, 33: S462.

[29]. Li Y., Qiu Y. and Zhu Y. Analysis and Application of EEG Signal Analysis Methods, pp. 49–51. Science Press, 2009.

[30]. Adeli H., Zhou Z., and Dadmehr N. “Analysis of EEG records in an epileptic patient using wavelet transform.” Journal of Neuroscience Methods. 2003, 123(1):69–87.

[31]. Nagelkerke, N. J. D. “A Note on a General Definition of the Coefficient of Determination”. Biometrika. 1991, 78(3): 691–692.

[32]. Chang C.-C. and Lin C.-J. “LIBSVM : a library for support vector machines”. ACM Transactions on Intelligent Systems and Technology. 2011, 2(27):1––27.

[33]. Peter L. et al. (2014) Leveraging Random Forests for Interactive Exploration of Large Histological Images. In: Golland P., Hata N., Barillot C., Hornegger J., Howe R. (eds) Medical Image Computing and Computer-Assisted Intervention – MICCAI 2014. MICCAI 2014. Lecture Notes in Computer Science, vol 8673. Springer, Cham.

[34]. Katharine M. Mullen. “Continuous Global Optimization in R”. Journal of Statistical Software. 2014,60(6).

[35]. Li H. Methods of statistical learning, pp. 67–71. QingHua University Press, 2012.

[36]. Mitchell, T. M. “Machine Learning”. China Machine Press. 2003, p.63.

